# The fall and rise of group B *Streptococcus* in dairy cattle: reintroduction due to human-to-cattle host jumps?

**DOI:** 10.1101/2021.04.21.440740

**Authors:** Chiara Crestani, Taya L Forde, Samantha J Lycett, Mark A Holmes, Charlotta Fasth, Karin Persson-Waller, Ruth N Zadoks

**Affiliations:** Institute of Biodiversity, Animal Health and Comparative Medicine, University of Glasgow, Garscube Campus, Glasgow G61 1QH, UK; The Roslin Institute, University of Edinburgh, Easter Bush Campus, Midlothian EH25 9RG, UK; Department of Veterinary Medicine, University of Cambridge, Madingley Rd, Cambridge CB3 0ES, UK; National Veterinary Institute (SVA), SE-751 89 Uppsala, Sweden; Moredun Research Institute, Pentlands Science Park, Bush Loan, Penicuik EH26 0PZ, UK; School of Veterinary Science, University of Sydney, Werombi Road, Camden, NSW 2570, Australia

**Keywords:** Streptococcus agalactiae, emergence, anthroponosis, reverse zoonosis, plasmid, host adaptation

## Abstract

Group B *Streptococcus* (GBS; *Streptococcus agalactiae*) is a major neonatal and opportunistic bacterial pathogen of humans and an important cause of mastitis in dairy cattle with significant impacts on food security. Following the introduction of mastitis control programs in the 1950s, GBS was nearly eradicated from the dairy industry in northern Europe, followed by re-emergence in the 21st century. Here, we sought to explain this re-emergence based on short and long read sequencing of historical (1953-1978; n = 44) and contemporary (1997-2012; n = 76) bovine GBS isolates. Our data show that a globally distributed bovine-associated lineage of GBS was commonly detected among historical isolates but never among contemporary isolates. By contrast, tetracycline resistance, which is present in all major GBS clones adapted to humans, was commonly and uniquely detected in contemporary bovine isolates. These observations provide evidence for strain replacement and suggest a human origin of newly emerged strains. Three novel GBS plasmids were identified, including two showing >98% homology with plasmids from *Streptococcus pyogenes* and *Streptococcus dysgalactiae* subsp. *equisimilis*, which co-exist with GBS in the human oropharynx. Our findings support introduction of GBS into the dairy population due to human to-cattle jumps on multiple occasions and demonstrate that reverse zoonotic transmission can erase successes of animal disease control campaigns.

**IMPACT STATEMENT:** Pathogens can jump between humans and animals. Animal domestication and intensification of livestock production systems have caused multiple human to animal spill-over events, sometimes with significant impact on animal health and food production. The most common production-limiting disease of dairy cattle is mastitis, inflammation of the mammary gland, which can be caused by group B *Streptococcus,* a common commensal and pathogen of humans. Using genomic data from historical and recent isolates, we show that re-emergence of this pathogen in the dairy industry in northern Europe is due to strains with genomic signatures of human host-adaptation, including antimicrobial resistance genes and plasmids. This shows how elimination of animal diseases may be hampered by humans serving as a reservoir of multi-host pathogens, and reverse zoonotic transmission.

**REPOSITORIES:** Reads for all isolates sequenced in this study have been submitted to the ENA Sequence Read Archive (SRA). SRA accession numbers are included in Table S1, supplementary material, available in the online version of this article.

## 1. Introduction

Group B *Streptococcus* (GBS), or *Streptococcus agalactiae*, is the leading cause of human neonatal meningitis in high income countries [1] and causes invasive and non-invasive disease in adults with or without underlying medical conditions [2,3]. GBS is also a commensal of the lower gastrointestinal and urogenital tract of men and women, with an estimated carriage prevalence of 20 to 30% [4,5]. Additional colonisation sites include the skin and oropharynx [5,6,7]. Many animal species can be infected with GBS, and major economic impacts are recognised in the global dairy and aquaculture industries. Emergence of GBS in animal production systems occurred concurrently with changes in husbandry practices, such as use of milking machines, or the intensification of commercial aquaculture [3,8,9].

In the 1950s and 1960s, mastitis control programs were implemented to limit the impact of GBS on milk production. Such programs focused on identification and antimicrobial treatment of infected cattle and prevention of GBS transmission during milking, and led to near-elimination of bovine GBS in Canada [10], the UK [11] and northern Europe [12,13,14], with elimination (“reduction to zero of the incidence of disease or infection in a defined geographical area” [15]) achieved by most farms in those areas. The success of GBS mastitis control programs, which predate genetic typing of bacterial isolates by several decades, was attributed to the perception that GBS is an “obligate intramammary pathogen of dairy cattle” [8], despite its prevalence in humans. In the UK [16], the USA [9], and Portugal [17], a single bovine-adapted lineage of GBS, clonal complex (CC) 61, predominates in cattle. This observation, combined with the absence of CC61 among human GBS collections, has fuelled the perception that this GBS lineage is bovine-specific citeRichards2019, Bisharat, Almeida.

In recent years, re-emergence of GBS in dairy herds has been documented in several Nordic countries [2,8,18]. This phenomenon is partly attributed to changes in dairy production systems, including herd size, ownership structure and management practices [8,18]. It is not clear, however, why approaches that were adequate for control of GBS in other decades or countries would fail in northern Europe, unless pathogen evolution has changed the paradigm on which these programs were built, necessitating the use of additional or alternative approaches. Such insight is fundamental to the development of new animal disease control strategies, especially those that do not rely on routine use of antimicrobials.

We hypothesised that the bovine-adapted GBS lineage CC61 was eliminated through dedicated mastitis control programs, with subsequent emergence of GBS from other lineages, possibly as a result of host-species jumping, as described for *Staphylococcus aureus* mastitis in cattle [19] and suggested for GBS in fishes [3]. To test this, we investigated GBS isolates collected from bovine milk in Sweden over six decades, focusing on shifts in population composition, and the detection of genetic markers of host adaptation that might provide insight into a potential reverse zoonotic origin of newly emerged GBS lineages in cattle.

## 2. Methods

### Bacterial Isolates

Historical (1953-1978; n = 45) and contemporary (1997-2012; n = 77) bovine GBS isolates were obtained from the National Veterinary Institute (SVA; Table S1). No isolates were available from 1979 through 1996 (inclusive). Isolates originated from bovine milk samples from 107 farms and had been submitted to SVA for diagnostic testing. In Europe, GBS isolates from a dairy farm generally belong to a single strain or sequence type (ST) [18,20]. Therefore, one isolate per farm per year was selected for sequencing, with one exception (Table S1). Archived isolates were plated on sheep blood agar (E&O Laboratories) and grown overnight at 37°C to confirm viability and purity. An isolated colony was inoculated into Todd Hewitt broth (Oxoid-Thermo Fisher Scientific) and incubated aerobically at 37°C for 24 hours to prepare a bacterial suspension for subsequent DNA extraction.

### Short Read Sequencing

DNA was extracted with the GenElute Bacterial Genomic DNA Kit (Sigma-Aldrich) as per the manufacturer’s instructions. Library preparation was carried out with the Nextera XT DNA Sample Preparation Kit and MiSeq Reagent Kit V2 Library Preparation Kit (Illumina Inc.). DNA was sequenced with Illumina MiSeq technology. Paired-end raw reads were trimmed and filtered with ConDeTri v2.3 to remove low-quality bases and PCR duplicates [21] and *de novo* assembly was performed with SPAdes v3.11.1 [22] (the complete assembly pipeline can be found in the supplementary material).

Quality control of the assemblies generated from Illumina data (n=122) was carried out with QUAST v5.0.2 [23], with 2603V/R (human GBS, ST110, CC19, accession NC_004116) as the reference genome. Results for the total length of the genome, total number of contigs, N50 and GC content were plotted with the Python Seaborn library [24] (Fig. S1) and low-quality genomes were identified with a custom-built bash pipeline (see supplementary material). Dataset mean values for genome length, total number of contigs and N50 were 2,126,345 bp, 58 and 492,052 bp, respectively. Two genome assemblies were excluded from subsequent analyses: the first had a high GC content compared to the dataset average (isolate GC = 36.92%, dataset mean plus twice standard deviation = 35.43% ± 0.32). The sequence was checked with KmerFinder v3.1 [25] and was identified as belonging to a different bacterial species, *Enterococcus thailandicus*. The second genome had low quality scores for total number of contigs (n = 1,837), N50 (1,992 bp) and genome length (2,751,323 bp), which are indicative of possible contamination. Therefore, only 120 high-quality genome assemblies were selected for subsequent analyses. The distribution of contig numbers for the 120 high quality assemblies was bimodal (Fig. S2), with most of the more fragmented genomes belonging to bovine-adapted lineage CC61 (mean contig number = 112, compared to mean contig number = 35 for other genomes; Fig. S2B). Genome fragmentation was attributed to presence of a relatively high number of mobile genetic elements (MGE) and insertion sequences (IS) in this lineage [9,26].

### Long Read Sequencing

To obtain closed circular genomes, Oxford Nanopore MinION sequencing [27] was applied to a subset of isolates (n=22, Table S1). Within each lineage, isolates were selected to maximise the diversity in terms of ST, antimicrobial resistance determinants and presence/absence of integrative conjugative elements (ICE) based on analysis of the short read sequencing data. Two libraries, each consisting of 11 samples and a negative control, were prepared with the Rapid Barcoding Kit (SQK-RBK004 - Oxford Nanopore Technologies) and sequenced for 2 to 5 hours, generating an average of 1.73 Gb per run, with an estimated mean sequence coverage of 60x. Guppy v3.3.0 [28] was used for base calling and demultiplexing, and Unicycler v0.4.8 [29] was used to generate high-quality hybrid assemblies of raw Nanopore and Illumina data. Unicycler was able to resolve 20 complete genomes out of 22 generated using Oxford Nanopore MinION sequencing. Nineteen of these were generated with a hybrid Illumina-Nanopore reads assembly, and one (MRI Z2-182) using long-read-only assembly. The hybrid assembly for this genome generated eight contigs. We were not able to resolve closed genomes for isolates MRI Z2-332 and MRI Z2-340.

### Core Genome Analysis

A core genome alignment was obtained with Parsnp v1.2 [30]. RAxML-NG v0.9.0 [31] was used to infer a maximum-likelihood tree under a GTR+G model, which was inspected and annotated using iTOL [32]. Nucleotide sequences were annotated with Prokka v1.13.7 [33], and Roary v3.12.0 [34] was used to generate a pangenome. To investigate unresolved relationships between isolates that could be caused by recombination in the core genes, SplitsTree v4.15.1 [35] was used (Fig. S3).

MLST profiles were identified with SRST2 v0.2.0 [36] and capsular serotyping was conducted *in silico* following the method described by the Centers for Disease Control and Prevention (CDC) [37]. Briefly, BLASTn was used to search genome assemblies for the presence of serotype-specific short sequences extracted from the capsular serotype operon of selected reference genomes. With this approach, a perfect identity match is required for serotype VII and IX, whereas a minimum identity (ID) of 96% is suggested for serotypes Ia, Ib, and II through VI. We first validated this method on a database of publicly available GBS genomes [38], comprising human and animal sequences. Whole genome sequence (WGS) serotyping results matched perfectly with phenotypic typing methods. A lower ID threshold was observed for most serotype Ia strains in our study compared to the CDC study [37], as the majority of serotype Ia nucleotide sequences had a 94% ID match. Although most genomes had only one best match, two best matches were observed in a few cases. For these, the sequences were re-analysed using an *in silico* serotyping method that is based on the alignment of longer serotype-specific capsular operon sequences [39].

### Analysis of Accessory Genome Content

Antimicrobial resistance genes were detected with ResFinder v3.2 [40]. Presence of the lactose operon (Lac.2) [26,41,42], which is a marker of bovine host adaptation, was assessed with BLASTn (query coverage QC>90%, and ID 95%), searching for genotypes Lac.2a, Lac.2b and Lac.2c [41]. Detection of ICE Tn*916* and Tn*5801*, which carry the tetracycline resistance genes typical of human-associated GBS lineages [38], was also conducted with BLASTn searches (QC >80% and ID>95%), using reference sequence *S. agalactiae* 2603V/R, ICESag2603VR-1 (length = 18,031 bp) and *S. agalactiae* COH1, AAJR01000021.1 (selected region from 14,055 to 34,289; length = 20,235 bp), respectively. When *tet* (M)−positive sequences showed high identity (ID>95%) but only partial query coverage (QC<80%) with either of the two elements, an area of 20,000 bp surrounding *tet* (M) was manually selected and BLASTn was used to determine the ICE family with ICEfinder [43].

Lac.2 variants and ICE sequences were extracted from the genomes for phylogenetic analysis with ARIBA v2.14.4 [44] and manual curation when only a partial sequence was obtained because of divergence from the reference. Manual extraction of amino acid sequences was carried out from annotation files for the Lac.2 integrases genes, when possible, and for the *tet* (M) gene. Alignments of the nucleotide sequences of the ICE and the Lac.2 variants, and of the amino acid sequences of the *tet* (M) and the Lac.2 integrase genes, were carried out with MAFFT v7.407 [45] and Neighbor-Joining trees were built within Geneious software [46] with a Jukes Cantor model (default settings) (Fig. S4 and Fig. S5).

Figures were edited using Inkscape (http://www.inkscape.org).

## 3. Results and Discussion

### Eradication and Re-emergence of GBS is Associated with Strain Replacement

To test the hypothesis that the near-elimination and re-emergence of GBS in the Swedish dairy cattle population was associated with strain replacement, we inferred sequence types (ST) from the genomes of 120 GBS isolates from bovine milk, including 44 historical isolates collected from 1953 to 1978 and 76 contemporary isolates collected from 1997 to 2012. GBS detection in milk was exceedingly rare in the intervening period, and no stored isolates were available for typing. Bovine-adapted lineage CC61 was exclusively detected among historical isolates collected before 1970, as was minor lineage CC297. By contrast, two other lineages (CC1 and CC103/314) were only detected among contemporary isolates (Fig. 1). CC61, also referred to as CC61/67 or CC67, was first recognised in the UK [16] and is also widespread in cattle in the USA [9] and Portugal [17]. With the exception of three recent cases in China [47], CC61/67 has never been reported in people. Its absence from humans may be due to pseudogenisation of the operon that encodes the polysaccharide capsule, an important virulence factor in human but not bovine GBS infections [17]. Without alternative host species, elimination of CC61 from the cattle population would mean that no reservoir is left, precluding re-emergence and explaining its absence among contemporary isolates. Not much is known about the origin or fate of CC297, which is a rare type in humans as well as animals. Contemporary lineage CC1, by contrast, is common among human carriage and disease isolates, including in Sweden [2,48]. It has recently been recognised as a common cause of bovine mastitis in northern Europe [2,18,20] and elsewhere [6]. Contemporary lineage CC103/314 is recognised as a human pathogen in Asia, including Thailand [49], Taiwan [50], and China [51]. In cattle, it is found across multiple continents, with reports of CC103/314 as a common lineage among bovine isolates from China [52], Colombia [6], Denmark [20], Finland [2] and Sweden (Fig. 1). Re-emergence of pathogens may be due to cessation of control activities once near-elimination is achieved, with or without re-introduction of pathogens [15]. In Northern Europe, changes in animal husbandry and transmission patterns may have contributed to GBS re-emergence [2,18], and the lineage-replacement we describe here shows that re-introduction of GBS must also have occurred.

**Figure 1.**
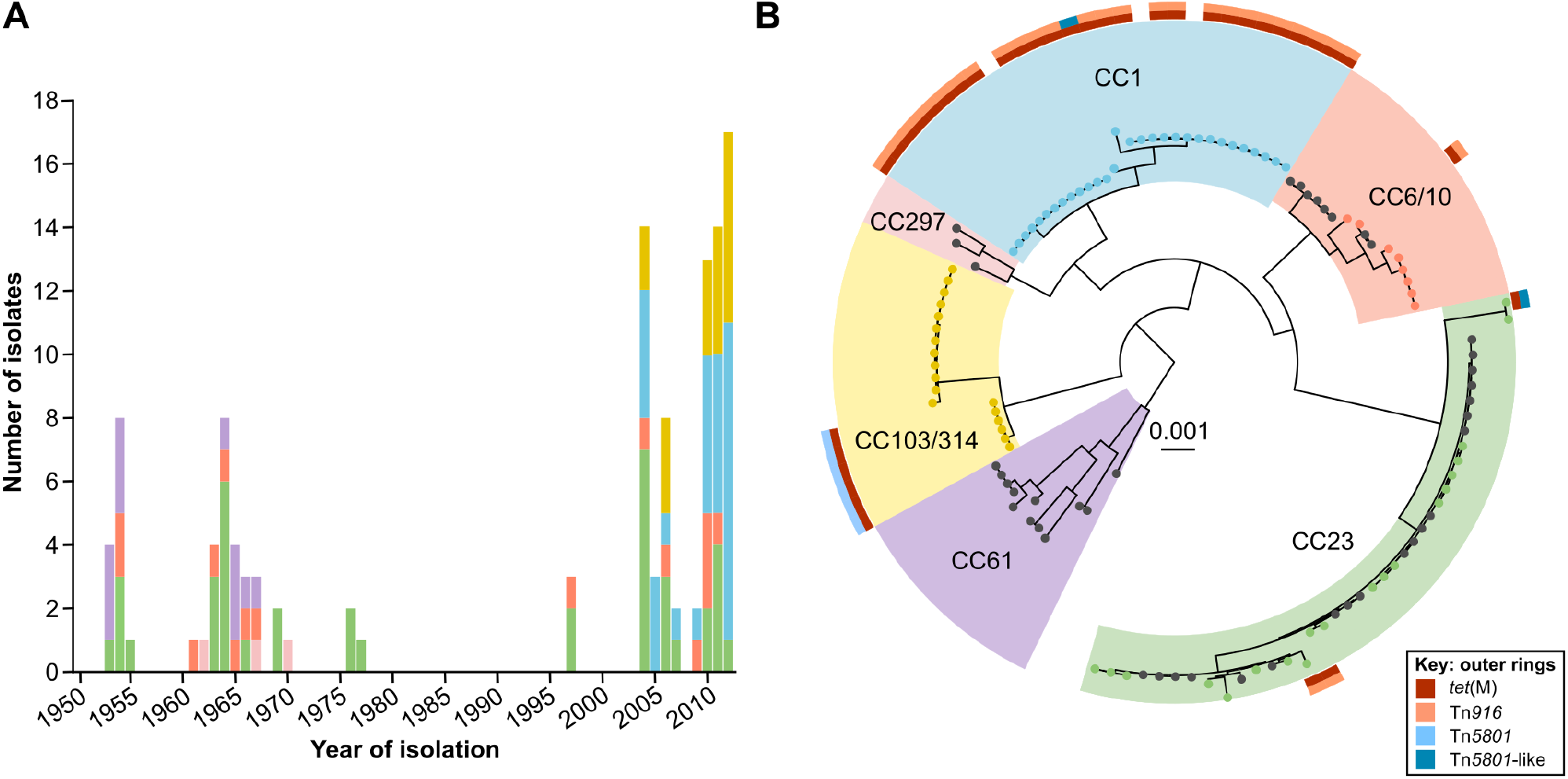
Control and re-emergence of group B *Streptococcus* (GBS) is associated with strain replacement. A) Frequency distribution of 120 GBS isolates from bovine milk, collected from 1953 to 1977 (n = 44) and 1997 to 2012 (n = 76), shows presence of six clonal complexes (CC; colour as indicated in panel B). Bovine-specific GBS lineage CC61 (purple) was last detected prior to 1970. Re-emergence of GBS in the Swedish dairy cattle population after a period of near-elimination is associated with CC1 (blue) and CC103/314 (yellow), which were first detected in 2004. Lineages CC23 (green) and CC6/10 (orange), which are also commonly found in other host species, were detected among historical and contemporary isolates. B) Tetracycline resistance, a marker of adaptation of GBS to humans, was detected exclusively among contemporary bovine GBS isolates. The core genome phylogeny of historical and contemporary isolates (black and coloured leaves, respectively) is shown, with clonal complex (CC) and presence of *tet* (M) and integrative conjugative elements (ICE), namely Tn*916*, Tn*5801* and Tn*5801*-like. *tet* (M) was carried by Tn*5801* in all ST314 isolates, and mostly by Tn*916* among CC1 isolates. One ST28 isolate from 1978 did not belong to any of the major clades and is not shown. Tree was rooted at midpoint.

### Host-adaptation Markers Suggest Human-to-Bovine Host Jumps

The majority of CC1 (27 of 30, 90%) and many CC103/314 (7 of 18, 39%) isolates carried the tetracycline resistance (TcR) gene *tet* (M), which was not detected in any historical isolates. TcR genes were carried by ICE Tn*916* or Tn*5801* or a Tn*5801*-like element (29, 6 and 2 of 37 *tet* (M) positive genomes, respectively; supporting material, Fig. S4). All bovine CC1 isolates belonged to serotype V, and TcR was predominantly associated with Tn*916* within this clade. TcR in CC103/314 was exclusively associated with Tn*5801* (Fig. 1). TcR is rare among bovine isolates but very common among human isolates [9]. In deed, the human GBS population is dominated by a few GBS lineages that expanded after acquisition of TcR [38]. We interpret the presence of TcR in newly emerged bovine GBS lineages as an indication that those lineages have a human origin. In human GBS, CC1 serotype V acquired Tn*916* with TcR around 1935 [38]. Tn*5801* carrying TcR was acquired by human GBS around 1920 for CC17 and around 1950 for CC23, with no year reported for CC10 [38]. Since their acquisition, TcR determinants have persisted in the human GBS population even in the absence of selective pressure, presumably as a result of low fitness cost [38]. In our study, other tetracycline resistance genes (*tet* (A) and *tet* (K)) were detected once and three times, respectively, macrolide and lincosamide resistance genes *erm*B and *lnu*A and amino-glycoside resistance gene *str* were detected once, and chloramphenicol resistance genes *cat* (pC221) and *lsa*C were detected 2 and 5 times, respectively (Table S1). The low prevalence of resistance reflects longstanding restrictive veterinary antimicrobial use policies in Sweden, and the use of narrow spectrum penicillin as drug of first choice for bovine GBS treatment [53].

### Bovine GBS Shares Plasmids with Human-pathogenic Group A *Streptococcus* and Group G Streptococcus

Plasmids are rarely reported in GBS [9], but using high-quality circularised hybrid assemblies of raw Nanopore and Illumina data obtained for a subset of 20 GBS isolates, we identified three plasmids and one integrative element among four complete hybrid assemblies, belonging to three different lineages (Fig. 2). Plasmid pZ2-265 (isolate MRI Z2-265, ST61, CC61, length = 3,617 bp, accession MW118669), was nearly identical (QC 100%, ID 99.28%) to plasmid pA996 from *Streptococcus pyogenes* or group A *Streptococcus*, GAS [54]. The plasmid was detected in five isolates belonging to CC61 (twice using hybrid assembly and three times using BLASTn on short-read assemblies; Table S1). It encodes a toxin/antitoxin system, comprising a toxin of the RelE/ParE superfamily, and a prevent-host-death antitoxin (*phd*) (Fig. 2A). The latter represses transcription of the toxin and prevents host death by binding and neutralising the toxin [55]. Plasmid pZ2-174 (isolate MRI Z2-174, ST314, CC103/314, length = 3,041 bp, accession MW118668) showed significant similarity with plasmid pW2580 (QC 99%, ID 98.85%) from another human-associated pyogenic streptococcal species, *Streptococcus dysgalactiae* subsp. *equisimilis* or group G *Streptococcus* (GGS). This plasmid encodes the dysgalactin gene (*dysA*) (Fig. 2B), a bacteriocin directed primarily against *S. pyogenes* [56] and its immunity factor (*dysI*) [57].

**Figure 2.**
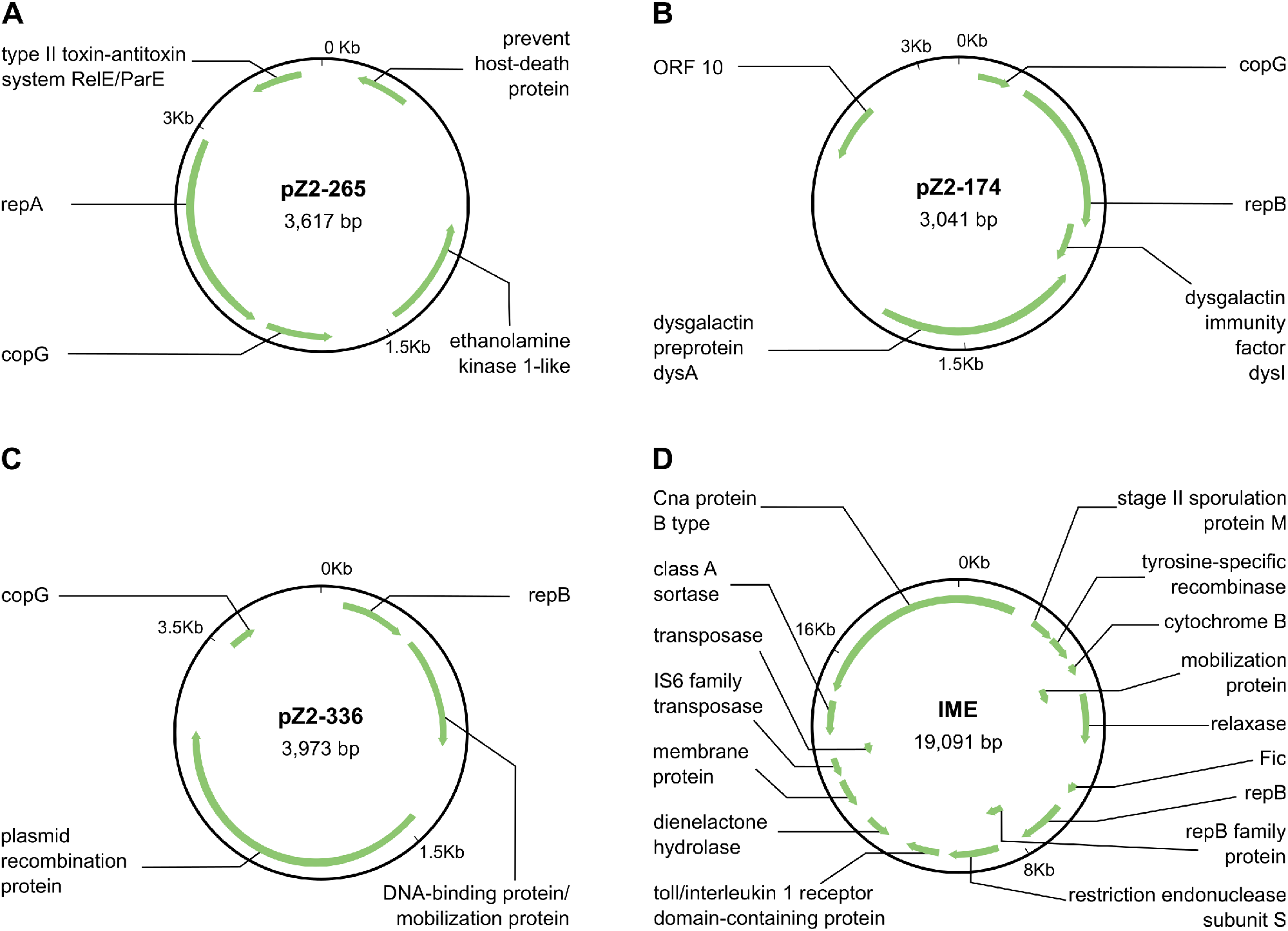
Hybrid Illumina-MinION assemblies of bovine group B *Streptococcus* (GBS) revealed the presence of plasmids and integrative mobilizable elements (IME). A) Plasmid pZ2-265 has 99.28% homology to plasmid pA996 (KC895877.1) from *Streptococcus pyogenes* or group A *Streptococcus*, GAS. B) Plasmid pZ2-174 shows 98.85% homology to pW2580 (AY907345.1) from *Streptococcus dysgalactiae* subsp. *equisimilis* or group G *Streptococcus*, GGS. C) pZ2-336 did not show significant similarity with known plasmids, whilst a second circular element in the same genome assembly (D) could either belong to a novel unclassified mobile genetic element family or be an IME.

GAS is uniquely associated with human hosts, and co-exists with GBS in the human oropharynx, as does GGS [7]. pW2580 may provide a survival advantage to GGS when it co-exists with GAS in the oropharynx. Exchange of plasmids or other mobile genetic elements between GAS, GBS and GGS is possible in this niche [7] and could potentially be followed by human to bovine transmission of GBS, as documented in epidemiological and evolutionary studies [9,58]. Finding two plasmids previously associated with other human *Streptococcus* species in bovine GBS isolates suggests that reverse zoonotic events (i.e. human-to-bovine spill-over) have occurred more than once. It is conceivable, albeit speculative, that early plasmid acquisition by GBS (pZ2-265 from GAS) and spill-over occurred prior to the expansion of CC61/67 in cattle, and that human GBS subsequently acquired other plasmids (pZ2-174 from GGS) and TcR prior to the recent expansion of CC1 and CC103/314 in cattle.

The third plasmid identified in this study, pZ2-336, (isolate MRI Z2-336, ST8, CC6/10, length = 3,973 bp, accession MW118670) did not show significant similarity with known plasmids. It encoded genes for plasmid mobilisation and recombination but no genes involved in bacterial protection or toxicity (Fig. 2C). In the same genome assembly, a second circular element was detected (length = 19,091 bp, accession MW118671), showing features of ICE or integrative mobilizable element (IME; tyrosine recombinase/integrase, relaxase), plasmids (plasmid mobilisation protein, plasmid replication initiation protein *repB*) and insertion sequences (IS; IS6 family transposase) (Fig. 2D). Additionally, it encoded genes with functions of cell adhesion (Cna protein B-type domain superfamily) and virulence factor expression (class A sortase). ICEFinder [43] identified a segment of this element (length = 11,068bp) as a putative IME. Hence, this newly described element could either belong to a novel unclassified family of mobile genetic elements (MGE) or it could be an actual IME.

None of the plasmids carried antimicrobial resistance genes.

### Multi-host Lineages of GBS Occur among Historic and Contemporary Bovine GBS Isolates

Two lineages, CC23 and CC6/10, were identified among both historical and contemporary bovine isolates (Fig. 1). CC23 is a common cause of bovine mastitis in northern Europe, whilst CC6/10 is less prevalent [2,20]. Both lineages affect humans and terrestrial and aquatic animals, including homeothermic species and poikilothermic species, e.g. seals and crocodiles, respectively, for CC23, or dolphins and fishes, respectively, for CC6/10, and are considered to be multi-host lineages [9,59,60]. Multiple serotypes are associated with both lineages [9]. For CC23 serotype Ia is primarily found in humans and serotype III in cattle [2,41]. In our study, isolates from CC23 mostly belonged to serotype III although a few serotype Ia isolates were detected in both eras. Four serotypes were identified in CC6/10 (Table S1). The detection of multihost lineages among historical and contemporary isolates could reflect ongoing low-level transmission in cattle during the interim period, as suggested by the dominance of serotype III in CC23. Alternatively or additionally, it could be due to sporadic reverse zoonotic transmission, as suggested in studies from Colombia [6], Denmark [8,41], and the USA [58], and compatible with occasional detection of CC23 isolates with the predominantly human-associated serotype Ia.

### Genome Plasticity Facilitates Host-adaptation

Based on analysis of 120 Illumina assemblies, the bovine GBS pangenome comprised 7,845 genes, of which the majority (80.3% or 6,297 genes) were accessory genes (989 shell genes and 5,308 cloud genes), with 17.8% core genes (present in all genomes) and a minority of soft core genes (present in 95 to 99% of genomes). The lactose operon Lac.2 was detected in almost all isolates in our study (Table S1). Lac.2 encodes the metabolism of lactose, or milk sugar, which constitutes a major adaptation of bovine GBS to the mammary gland, whereas it is largely absent from human GBS [2,26]. Integration sites for the lactose operon included the N-6 DNA methylase gene (N-6DNAM), *yxdL* (a multi-copy gene), *rbgA*, *lacD* from the Lac.1 operon [26], *gcvT* and hypothetical genes (Table S1), whereby each Lac.2 variant could be integrated at multiple sites, e.g. *lacD* or *rgbA* for Lac.2a, N-6DNAM or *yxdL* for Lac.2b and N-6DNAM or *gcvT* for Lac.2d positive isolates (Fig. 3). The latter is a new Lac.2 operon identified in this study that combines features of Lac.2a and Lac.2b (Fig. S5) [26,41]. Phylogenetic analysis showed that closely related Lac.2 sequences can belong to different variants (Fig. S5) and multiple Lac.2 variants were identified within most STs (Table S1). The heterogeneous distribution of the lactose operon and the diversity of integration sites illustrates the genome plasticity of GBS, which facilitates acquisition of accessory genome content and migration between host species [9].

**Figure 3.**
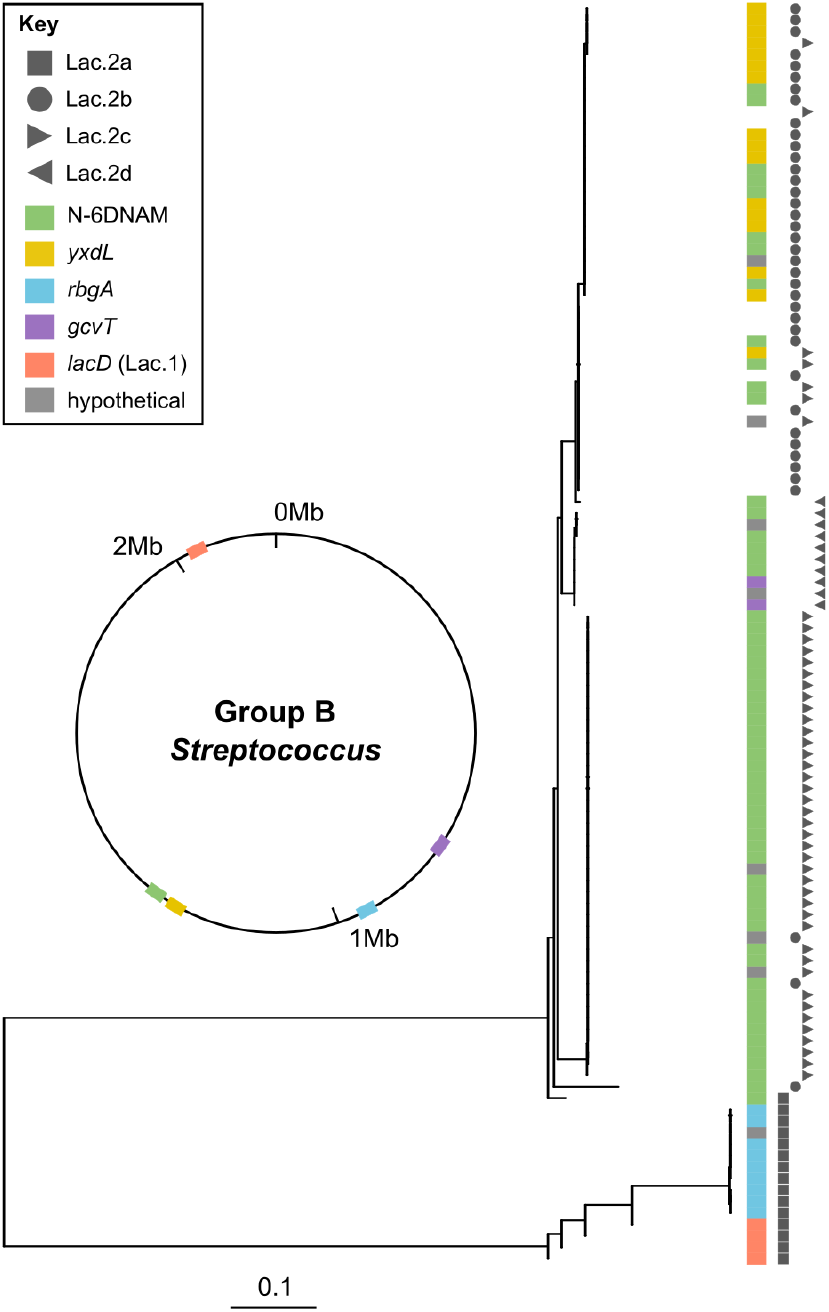
Phylogeny of the Lac.2 integrase amino acid sequences from bovine group B *Streptococcus* (GBS), with their insertion site (coloured strip) and Lac.2 variant (symbols), illustrating the plasticity of the accessory genome. Insertion sites have been mapped on an example GBS genome. Gene *yxdL* was found in multiple copies within the same genome, with Lac.2 detected next to the copy present in the region around 1.25 Mbp. When Lac.2 was found at the edge of a contig, it was not possible to determine the site of integration (n = 13, blank colour strip) and the integrase sequence (n = 10, not present in tree).

### Concluding Comments

Although often described as an obligate intramammary pathogen of dairy cattle in the veterinary literature, GBS is a multi-host pathogen and a host-species jumper with diverse habitats on and off-farm [2,9,18]. Evolutionary evidence shows that human-to-bovine jumps are twice as likely as migration in the opposite direction [9]. Here, we provide evidence that elimination of a major bovine-adapted lineage (CC61) in Swedish dairy cattle was followed by emergence of new lineages that carry evolutionary evidence of human origin in the form of TcR markers [38], suggesting introduction of human lineages into the cattle population through reverse zoonotic transmission. Subsequently, these new lineages likely established themselves in cattle with the acquisition of the lactose operon Lac.2 [26], which represents the most important marker of the bovine-specific GBS accessory genome known to date. This sequence of events is supported by the fact that TcR is largely retained even in the absence of selective pressure [38], such as in the Swedish dairy industry where antibiotic usage is low. The lactose operon does not appear to be retained outside of the bovine host [2,41]. Thus, TcR and Lac.2 provide historical, or long-term, and recent, or short-term, “records” of host adaptation, respectively.

Due to the unique historical nature of our isolate collection, direct comparison with genomic sequences of human isolates from the same area and era is not possible. Such comparisons, however, are not necessary for evolutionary analysis, whereby host species jumps have commonly been inferred based on sequence data of isolates derived from different host species without known interactions or epidemiological relatedness [19,61,62]. For the emergence of GBS in farmed species, several routes of transmission from humans to animals can be envisaged, including, in the case of fishes, the use of human waste for nutrient recycling and, in the case of cattle, the handling and milking of cows, which may lead to direct human-to-animal transmission [6,58,41]. Changes in animal husbandry systems combined with pathogen evolution are the likely explanation for the re-emergence of GBS, which has been observed in several countries in Europe [2,18,63].

Of the two emerging lineages in cattle, CC1 is known to co-circulate in the human and bovine populations in northern Europe [2]. By contrast, CC103/314 is common in dairy cattle on multiple continents but rare in humans, with the exception of Asia. Despite its low prevalence in humans, CC103 may have emerged in cattle due to a spill-over event, with subsequent amplification in modern dairy systems. There is precedent for such a chain of events, as there is reasonable evidence that GBS ST283, which is rare among human GBS isolates, emerged in aquaculture during its intensification in Asia as the result of spill-over from humans, with acquisition of fish-associated MGE facilitating this process [3,60]. Host switching exposes GBS to different selective pressures and sources of accessory genome content [9], including plasmids, as demonstrated for GBS, GAS and GGS in the human oropharynx [7], and other MGE, as demonstrated for the lactose operon in GBS, *Streptococcus uberis* and *Streptococcus dysgalactiae* subsp. *dysgalactiae* in the bovine udder [26]. We propose that the concept of “genetic species” and “ecological species”, as previously described for *Thermotoga* spp. also applies to streptococci (Fig. 4) [64]. As farming systems, host contact structures, and selective pressures change, new strains and transmission routes of GBS may continue to emerge through zoonotic and reverse zoonotic transmission, potentially erasing the success of decades of disease control efforts or creating new threats to animal and public health. Control of GBS and other multi-host pathogens will require ongoing monitoring of pathogen diversity across host species and adaptive management in response to changing selective pressures and emergence of new pathogen strains.

**Figure 4.**
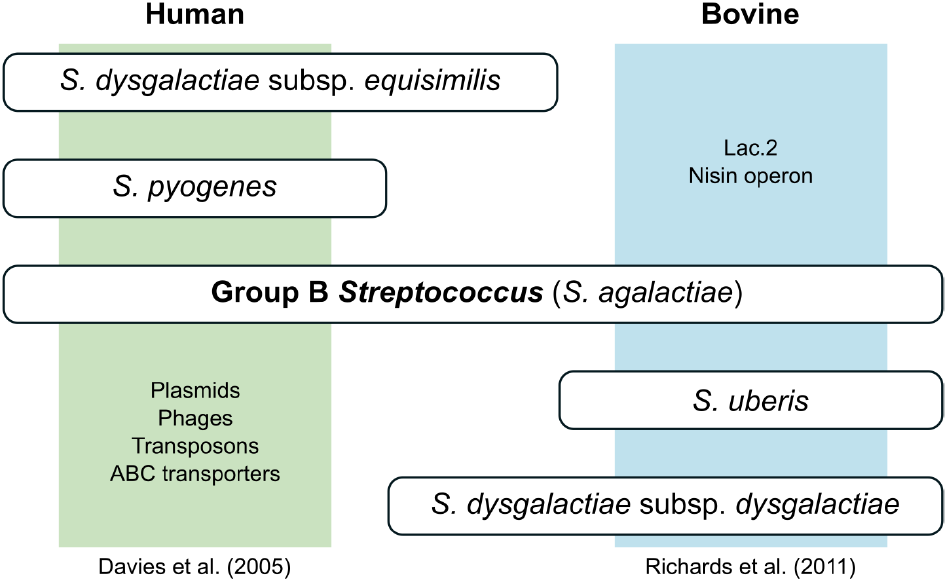
Schematic representation of the relationship between bacterial species and host species. The bacterial core genome defines bacterial species identity whereas the bacterial accessory genome drives and is driven by the host species inhabited by the bacteria. Acquisition of host-associated mobile genetic elements through lateral transfer between pyogenic streptococcal species in one host, as described for phages, transposons, and transporters in humans [7], and for lactose and nisin operons in cattle [26], followed by transmission between host species, may explain the detection of human-associated accessory genome content in bovine group B *Streptococcus* as observed in this study for tetracycline resistance and plasmids.

## Supporting information

Supportin_information

## 4. Author statements

### Authors and contributors

Conceptualisation: KPW and RNZ. Isolate and data curation: KPW, CF, MAH and RNZ. Formal analysis, investigation, methodology, visualisation: CC. Supervision: TF, SL and RNZ. Writing original draft: CC and RNZ. Writing review & editing: CC, RNZ, TF, MAH, SL, KPW.

## Acknowledgements

We thank Ian Heron and John Bell from the Moredun Research Institute for laboratory assistance, Kirstyn Brunker from the University of Glasgow for helping with the setup of Oxford Nanopore sequencing in the OHRBID laboratory, and Dr Ennio Lavagnini from the University of Cambridge for advice on development of the genome assembly pipeline for short paired-end reads.

## 5. Data summary

The authors confirm all supporting data, code and protocols have been provided within the article or through supplementary data files.

The scripts used for raw read assembly and genome quality control can be found in the supplementary file, available in the online version of this article.

Other external data used:

(1) 2603V/R, GenBank accession NC_004116
(2) Reference sequence Tn*916* from *S. agalactiae* 2603V/R, ICESag2603VR-1
(3) Reference sequence Tn*5801* from *S. agalactiae* COH1, AAJR01000021.1
(4) Plasmid pA996, KC895877.1 (5) Plasmid pW2580, AY907345.1

