## Supplementary material for "The fall and rise of group B *Streptococcus* in dairy cattle: reintroduction due to human-to-cattle host jumps?": Supportin_information

### Dataset 1

**Table S1** (Following page) Metadata and results of genomic analyses for 120 high-quality group B *Streptococcus* isolates from dairy cattle in Sweden. Columns show sample ID, accession number, year of isolation, serotype (from WGS), clonal complex (CC), sequence type (ST), antimicrobial resistance genes (AMR), presence and type of integrative conjugative elements (ICE) and lactose operon (Lac.2), Lac.2 insertion site, presence and type of plasmids, long-read MinION sequencing and farm of origin.

| SAMPLE | ACCESSION | YEAR | SEROTYPE | CC | ST | AMR | ICE | LAC.2 | LAC.2 INSERTION SITE | PLASMIDS | MINION | FARM |
|---|---|---|---|---|---|---|---|---|---|---|---|---|
| MRIZ2-138 | ERS1796664 | 2010 | NT | 6/10 | 10 | | | Lac.2a | rbgA | | | 74 |
| MRIZ2-139 | ERS1796665 | 2010 | III | 23 | 722 | *tet*(M) | Tn*916* | Lac.2a, Lac.2c | N-6DNAM, lacD | | | 29 |
| MRIZ2-140 | ERS1796666 | 2010 | IV | 1 | 196 | *tet*(M) | Tn*916* | Lac.2b | na | | | 47 |
| MRIZ2-141 | ERS1796672 | 2010 | V | 1 | 1 | *tet*(M) | Tn*916* | Lac.2b | N-6DNAM | | | 10 |
| MRIZ2-142 | ERS1796673 | 2010 | Ia | 103/314 | 103 | | | Lac.2b | yxdL | | | 106 |
| MRIZ2-143 | ERS1796674 | 2010 | Ia | 6/10 | 724 | | | Lac.2c | hypothetical protein | | | 19 |
| MRIZ2-144 | ERS1796680 | 2010 | Ia | 103/314 | 314 | *tet*(M) | Tn*5801* | Lac.2b | yxdL | | | 59 |
| MRIZ2-145 | ERS1796681 | 2010 | II | 6/10 | 10 | | | Lac.2a | rbgA | | | 49 |
| MRIZ2-146 | ERS1796682 | 2010 | V | 1 | 1 | *tet*(M) | Tn*916* | Lac.2b | hypothetical protein | | | 75 |
| MRIZ2-147 | ERS1796688 | 2010 | III | 23 | 723 | lsaC | | Lac.2c | N-6DNAM | | | 45 |
| MRIZ2-148 | ERS1796689 | 2010 | IV | 1 | 1384 | *tet*(M) | Tn*916* | Lac.2b | na | | | 24 |
| MRIZ2-149 | ERS1796690 | 2010 | IV | 1 | 196 | | | Lac.2b | na | | yes | 13 |
| MRIZ2-150 | ERS1796696 | 2010 | Ia | 103/314 | 103 | | | Lac.2b | yxdL | | | 38 |
| MRIZ2-152 | ERS1796697 | 2011 | NT | 1 | 1387 | *tet*(M) | Tn*5801*-like | Lac.2b | N-6DNAM | | | 23 |
| MRIZ2-153 | ERS1796698 | 2011 | Ia | 103/314 | 103 | | | Lac.2b | yxdL | | | 79 |
| MRIZ2-154 | ERS1796704 | 2011 | V | 1 | 1 | *tet*(M) | Tn*916* | Lac.2d | N-6DNAM | | | 50 |
| MRIZ2-155 | ERS1796705 | 2011 | III | 23 | 23 | | | Lac.2c | N-6DNAM | | | 105 |
| MRIZ2-156 | ERS1796706 | 2011 | IV | 1 | 726 | *tet*(M) | Tn*916* | Lac.2b | na | | | 72 |
| MRIZ2-157 | ERS1796712 | 2011 | Ia | 103/314 | 727 | *tet*(M) | Tn*5801* | Lac.2c | yxdL | | | 20 |
| MRIZ2-158 | ERS1796713 | 2011 | Ia | 23 | 23 | | | Lac.2d | hypothetical protein | | yes | 63 |
| MRIZ2-159 | ERS1796720 | 2011 | Ia | 103/314 | 103 | | | Lac.2a | rbgA | | | 67 |
| MRIZ2-160 | ERS1796721 | 2011 | III | 23 | 23 | | | Lac.2c | N-6DNAM | | | 39 |
| MRIZ2-161 | ERS1796722 | 2011 | II | 6/10 | 10 | | | Lac.2c | N-6DNAM | | | 5 |
| MRIZ2-162 | ERS1796728 | 2011 | NT | 23 | 23 | | | Lac.2c | N-6DNAM | | | 26 |
| MRIZ2-163 | ERS1796729 | 2011 | V | 1 | 1 | *tet*(M) | Tn*916* | Lac.2c | N-6DNAM | | | 15 |
| MRIZ2-164 | ERS1796730 | 2011 | V | 1 | 1 | *tet*(M) | Tn*916* | Lac.2b | N-6DNAM | | | 37 |
| MRIZ2-165 | ERS1796752 | 2011 | Ia | 103/314 | 103 | | | Lac.2b | yxdL | | | 85 |
| MRIZ2-166 | ERS1796753 | 2012 | V | 1 | 1 | *tet*(M) | Tn*916* | Lac.2b | N-6DNAM | | | 65 |
| MRIZ2-167 | ERS1796740 | 2012 | V | 1 | 1 | *tet*(M) | Tn*916* | Lac.2a | rbgA | | | 56 |
| MRIZ2-168 | ERS1796741 | 2012 | Ia | 103/314 | 314 | *tet*(M) | Tn*5801* | Lac.2b | yxdL | | | 48 |
| MRIZ2-169 | ERS1796748 | 2012 | IV | 1 | 1384 | *tet*(M) | Tn*916* | Lac.2b | na | | | 64 |
| MRIZ2-170 | ERS1796749 | 2012 | Ia | 103/314 | 103 | | | Lac.2b | yxdL | | | 86 |
| MRIZ2-171 | ERS1796569 | 2012 | Ia | 103/314 | 103 | | | Lac.2b | yxdL | | | 22 |
| MRIZ2-172 | ERS1796570 | 2012 | IV | 1 | 196 | *tet*(M) | Tn*916* | | | | yes | 104 |
| MRIZ2-173 | ERS1796571 | 2012 | Ia | 103/314 | 314 | *tet*(M) | Tn*5801* | Lac.2b | yxdL | | | 11 |
| MRIZ2-174 | ERS1796572 | 2012 | Ia | 103/314 | 728 | *tet*(M) | Tn*5801* | Lac.2b | yxdL | pZ2-174 | yes | 108 |
| MRIZ2-175 | ERS1796573 | 2012 | Ia | 103/314 | 103 | | | Lac.2b | yxdL | | | 52 |
| MRIZ2-177 | ERS1796574 | 2012 | V | 1 | 1 | *tet*(M) | Tn*916* | Lac.2c | N-6DNAM | | | 9 |
| MRIZ2-178 | ERS1796575 | 2012 | IV | 1 | 726 | *tet*(A), *tet*(M) | Tn*916* | Lac.2b | na | | | 110 |
| MRIZ2-179 | ERS1796576 | 2012 | V | 1 | 1 | | | Lac.2b | N-6DNAM | | yes | 90 |
| MRIZ2-180 | ERS1796577 | 2012 | IV | 1 | 196 | *tet*(M) | Tn*916* | Lac.2b | na | | | 81 |
| MRIZ2-181 | ERS1796579 | 2012 | IV | 1 | 726 | *tet*(M) | Tn*916* | Lac.2b | na | | | 73 |
| MRIZ2-182 | ERS1796580 | 2012 | Ia | 23 | 23 | *tet*(M) | Tn*5801*-like | Lac.2b | N-6DNAM | | yes | 18 |
| MRIZ2-183 | ERS1796581 | 2012 | V | 1 | 1 | *tet*(M) | Tn*916* | Lac.2a | rbgA | | | 41 |
| MRIZ2-261 | ERS1796582 | 1953 | III | 61 | 1386 | | | Lac.2a | na | | | 35 |
| MRIZ2-262 | ERS1796583 | 1953 | III | 61 | 1386 | | | Lac.2a | na | | | 34 |

**Table S1 continued from previous page**

| SAMPLE | ACCESSION | YEAR | SEROTYPE | CC | ST | AMR | ICE | LAC.2 | LAC.2 INSERTION SITE | PLASMIDS | MINION | FARM |
|---|---|---|---|---|---|---|---|---|---|---|---|---|
| MRIZ2-263 | ERS1796585 | 1953 | III | 61 | 1386 | | | Lac.2a | na | | | 62 |
| MRIZ2-264 | ERS1796586 | 1953 | III | 23 | 23 | | | Lac.2c | N-6DNAM | | | 27 |
| MRIZ2-265 | ERS1796587 | 1954 | III | 61 | 1516 | | | Lac.2a | na | pZ2-265 | yes | 111 |
| MRIZ2-266 | ERS1796589 | 1954 | III | 23 | 23 | | | Lac.2c | N-6DNAM | | | 8 |
| MRIZ2-267 | ERS1796590 | 1954 | II | 61 | 61 | | | Lac.2a | na | pZ2-265 | | 7 |
| MRIZ2-269 | ERS1796591 | 1954 | Ib | 6/10 | 6 | | | Lac.2a | rbgA | | | 34 |
| MRIZ2-270 | ERS1796592 | 1954 | Ib | 6/10 | 1512 | | | Lac.2a | rbgA | | yes | 71 |
| MRIZ2-271 | ERS1796593 | 1954 | Ia | 23 | 23 | | | Lac.2c | N-6DNAM | | | 102 |
| MRIZ2-272 | ERS1796594 | 1954 | Ia | 23 | 23 | | | Lac.2c | N-6DNAM | | | 44 |
| MRIZ2-273 | ERS1796595 | 1955 | III | 23 | 23 | | | Lac.2c | N-6DNAM | | | 28 |
| MRIZ2-274 | ERS1796596 | 1961 | Ib | 6/10 | 1513 | | | Lac.2d | N-6DNAM | | | 98 |
| MRIZ2-275 | ERS1796597 | 1962 | II | 297 | 1510 | lsaC | | Lac.2d | hypothetical protein | | | 70 |
| MRIZ2-276 | ERS1796598 | 1963 | III | 23 | 23 | | | Lac.2c | hypothetical protein | | | 3 |
| MRIZ2-277 | ERS1796599 | 1963 | Ib | 6/10 | 6 | | | Lac.2c | N-6DNAM | | | 70 |
| MRIZ2-278 | ERS1796600 | 1963 | III | 23 | 23 | | | Lac.2c | N-6DNAM | | | 88 |
| MRIZ2-279 | ERS1796601 | 1963 | III | 23 | 23 | | | Lac.2c | N-6DNAM | | | 99 |
| MRIZ2-280 | ERS1796602 | 1964 | Ia | 23 | 23 | | | Lac.2c | N-6DNAM | | | 4 |
| MRIZ2-281 | ERS1796603 | 1964 | III | 23 | 23 | | | Lac.2c | N-6DNAM | | | 31 |
| MRIZ2-282 | ERS1796604 | 1964 | III | 23 | 23 | | | Lac.2c | N-6DNAM | | | 89 |
| MRIZ2-283 | ERS1796605 | 1964 | III | 23 | 23 | | | Lac.2c | N-6DNAM | | | 6 |
| MRIZ2-284 | ERS1796606 | 1964 | III | 23 | 23 | | | Lac.2c | N-6DNAM | | | 53 |
| MRIZ2-285 | ERS1796607 | 1964 | III | 23 | 23 | | | Lac.2c | N-6DNAM | | | 82 |
| MRIZ2-286 | ERS1796609 | 1964 | III | 61 | 1386 | | | Lac.2a | na | | yes | 84 |
| MRIZ2-287 | ERS1796610 | 1964 | Ib | 6/10 | 12 | | | Lac.2c | N-6DNAM | | | 1 |
| MRIZ2-288 | ERS1796611 | 1965 | Ib | 6/10 | 6 | | | Lac.2d | N-6DNAM | | | 84 |
| MRIZ2-289 | ERS1796612 | 1965 | III | 61 | 1394 | | | Lac.2a | na | pZ2-265 | | 61 |
| MRIZ2-290 | ERS1796614 | 1965 | II | 61 | 1515 | | | Lac.2a | na | pZ2-265 | | 36 |
| MRIZ2-291 | ERS1796616 | 1965 | II | 61 | 1392 | | | Lac.2d | gcvT | | | 54 |
| MRIZ2-293 | ERS1796617 | 1966 | Ib | 6/10 | 12 | | | Lac.2c | N-6DNAM | | | 1 |
| MRIZ2-294 | ERS1796618 | 1966 | Ib | 61 | 1393 | | | Lac.2d | N-6DNAM | | | 6 |
| MRIZ2-295 | ERS1796619 | 1966 | III | 23 | 23 | | | Lac.2c | N-6DNAM | | | 93 |
| MRIZ2-296 | ERS1796620 | 1954 | II | 61 | 1392 | | | Lac.2d | gcvT | | | 76 |
| MRIZ2-297 | ERS1796621 | 1967 | Ia | 297 | 1511 | lsaC | | Lac.2a | lacD | | | 95 |
| MRIZ2-298 | ERS1796622 | 1967 | Ib | 6/10 | 6 | | | Lac.2a | na | | | 42 |
| MRIZ2-299 | ERS1796623 | 1967 | II | 61 | 1514 | | | Lac.2a | lacD | pZ2-265 | yes | 33 |
| MRIZ2-301 | ERS1796624 | 1969 | III | 23 | 23 | | | Lac.2c | N-6DNAM | | | 94 |
| MRIZ2-302 | ERS1796625 | 1969 | III | 23 | 23 | | | Lac.2c | N-6DNAM | | | 1 |
| MRIZ2-304 | ERS1796626 | 1970 | Ia | 297 | 297 | lsaC | | Lac.2a | lacD | | yes | 103 |
| MRIZ2-305 | ERS1796627 | 1976 | III | 23 | 23 | | | Lac.2c | N-6DNAM | | | 101 |
| MRIZ2-306 | ERS1796628 | 1976 | III | 23 | 1509 | | | Lac.2c | N-6DNAM | | | 96 |
| MRIZ2-307 | ERS1796629 | 1977 | III | 23 | 23 | | | Lac.2c | N-6DNAM | | yes | 109 |
| MRIZ2-308 | ERS1796630 | 1978 | Ia | 28 | 28 | | | Lac.2a | na | | | unknown |
| MRIZ2-311 | ERS1796631 | 1997 | III | 6/10 | 10 | | | Lac.2a | rbgA | | | 46 |
| MRIZ2-312 | ERS1796632 | 1997 | III | 23 | 23 | lsaC | | Lac.2c | N-6DNAM | | | 80 |
| MRIZ2-313 | ERS1796633 | 1997 | III | 23 | 23 | | | Lac.2c | N-6DNAM | | | 21 |
| MRIZ2-314 | ERS1796634 | 2004 | III | 23 | 23 | *tet*(K) | | Lac.2c | N-6DNAM | | | 17 |

**Table S1 continued from previous page**

| SAMPLE | ACCESSION | YEAR | SEROTYPE | CC | ST | AMR | ICE | LAC.2 | LAC.2 INSERTION SITE | PLASMIDS | MINION | FARM |
|---|---|---|---|---|---|---|---|---|---|---|---|---|
| MRIZ2-315 | ERS1796635 | 2004 | III | 23 | 23 | | | Lac.2b | N-6DNAM | | | 21 |
| MRIZ2-316 | ERS1796636 | 2004 | Ia | 23 | 1507 | | | Lac.2c | hypothetical protein | | | 57 |
| MRIZ2-317 | ERS1796637 | 2004 | Ia | 103/314 | 103 | | | Lac.2b | yxdL | | | 69 |
| MRIZ2-318 | ERS1796638 | 2004 | V | 1 | 1 | *tet*(M) | Tn*916* | Lac.2d | N-6DNAM | | yes | 16 |
| MRIZ2-319 | ERS1796639 | 2004 | V | 1 | 1517 | *tet*(M) | Tn*916* | Lac.2b | N-6DNAM | | | 107 |
| MRIZ2-320 | ERS1796640 | 2004 | III | 23 | 23 | lnuA | | Lac.2c | N-6DNAM | | | 107 |
| MRIZ2-321 | ERS1796641 | 2004 | IV | 1 | 196 | *tet*(M) | Tn*916* | Lac.2b | na | | | 78 |
| MRIZ2-322 | ERS1796642 | 2004 | Ia | 103/314 | 103 | | | Lac.2a | hypothetical protein | | yes | 68 |
| MRIZ2-323 | ERS1796643 | 2004 | IV | 1 | 1384 | *tet*(M) | Tn*916* | Lac.2b | na | | | 25 |
| MRIZ2-324 | ERS1796644 | 2004 | III | 23 | 23 | | | Lac.2c | N-6DNAM | | | 40 |
| MRIZ2-325 | ERS1796645 | 2004 | III | 23 | 23 | | | Lac.2c | N-6DNAM | | | 87 |
| MRIZ2-326 | ERS1796646 | 2004 | III | 23 | 23 | | | Lac.2c | N-6DNAM | | | 54 |
| MRIZ2-327 | ERS1796647 | 2004 | II | 6/10 | 10 | | | Lac.2a | rbgA | | | 46 |
| MRIZ2-328 | ERS1796648 | 2005 | V | 1 | 1 | | | Lac.2b | N-6DNAM | | yes | 58 |
| MRIZ2-329 | ERS1796649 | 2005 | V | 1 | 1 | *tet*(M), str | Tn*916* | Lac.2b | N-6DNAM | | yes | 14 |
| MRIZ2-330 | ERS1796650 | 2005 | IV | 1 | 732 | *tet*(M) | Tn*916* | Lac.2b | na | | | 60 |
| MRIZ2-331 | ERS1796651 | 2006 | V | 1 | 1 | *tet*(M) | Tn*916* | Lac.2b | N-6DNAM | | | 77 |
| MRIZ2-332 | ERS1796652 | 2006 | V | 23 | 1385 | *tet*(M) | Tn*916* | Lac.2b, Lac.2c | hypothetical protein, na | | yes | 83 |
| MRIZ2-333 | ERS1796653 | 2006 | Ia | 103/314 | 103 | | | Lac.2c | yxdL | | | 55 |
| MRIZ2-334 | ERS1796654 | 2006 | III | 23 | 1508 | | | Lac.2c | N-6DNAM | | | 32 |
| MRIZ2-335 | ERS1796655 | 2006 | Ia | 103/314 | 314 | *tet*(M), cat(pC221) | Tn*5801* | Lac.2b | yxdL | | yes | 12 |
| MRIZ2-336 | ERS1796657 | 2006 | Ib | 6/10 | 8 | *tet*(M), cat(pC221) | Tn*916* | Lac.2a | N-6DNAM | pZ2-336, IME | yes | 30 |
| MRIZ2-337 | ERS1796658 | 2006 | III | 23 | 23 | | | Lac.2c | N-6DNAM | | | 43 |
| MRIZ2-338 | ERS1796659 | 2006 | Ia | 103/314 | 103 | *tet*(M), *tet*(K) | | Lac.2b | yxdL | | yes | 97 |
| MRIZ2-339 | ERS1796660 | 2007 | IV | 1 | 1384 | *tet*(M) | Tn*916* | Lac.2b | na | | | 25 |
| MRIZ2-340 | ERS1796661 | 2007 | III | 23 | 23 | *tet*(K) | | Lac.2c | N-6DNAM | | yes | 107 |
| MRIZ2-342 | ERS1796662 | 2009 | II | 6/10 | 10 | | | | | | yes | 46 |
| MRIZ2-343 | ERS1796663 | 2009 | V | 1 | 1 | *tet*(M), ermB | Tn*916* | Lac.2d | N-6DNAM | | | 66 |

#### Genome Assembly Pipeline for Short Paired-End Reads

This pipeline was designed to automate the trimming, filtering and assembly of Illumina paired-end reads. The script concatenates three sub-scripts, two of which are part of the ConDeTri suite and one of which is SPAdes assembler. This script was created with the contribution of Dr Ennio Lavagnini, University of Cambridge.

---

```
#!/bin/sh

# Create a list with input files (which are zipped fastq files) for the loop. The list of
    files is made only with read 1 of each pair of reads. Later the sed command will
    couple read 1 and read 2.

ls *1.fastq.gz > list

# Create the work folder (where the files will be processed) and the output folder (where
    the files will be stored).

mkdir Work_folder
mkdir Output
mkdir Output/Contigs
mkdir Output/Scaffolds
mkdir Output/Trim_reads

# This is the main loop of the script and it will move each couple of paired end reads in
    the Work_folder before running three different scripts (for trimming, filtering and
    assembling).

while read fast;
do

# The sed command is used to name the variables.

varzip1=$(echo $fast)
varzip2=$(echo $fast | sed s/_1.fas/_2.fas/)

# Unzipping the couple of files of interest (read 1 and read 2).

    gzip -d $varzip1
    gzip -d $varzip2

# Define 2 new variables for unzipped files. They are var1 = read 1 and var2 = read 2.

var1=$(echo $varzip1 | sed s/_1.fastq.gz/_1.fastq/ )
var2=$(echo $varzip2 | sed s/_2.fastq.gz/_2.fastq/ )

# Move unzipped files into Work_folder and define new variable "pref" that will be used in
    the scripts. The prefix variable will be the ID of the sequence, without any extension.

mv $var1 Work_folder && mv $var2 Work_folder
cd Work_folder
pref=$(echo $var2 | sed s/_2.fastq/""/)

# First script for trimming the reads (ConDeTri).
echo ""
echo "======Starting ConDeTri for $pref..."
echo ""

perl $HOME/anaconda3/envs/assembly/bin/condetri.pl -fastq1=$var1 -fastq2=$var2
    -prefix=$pref -hq=25 -lq=10 -frac=0.8 -minlen=50 -mh=5 -ml=1 -sc=33

echo ""
echo "======ConDeTri finished for $pref!"
echo ""
```

```
# Remove useless files.

rm *stats ; rm *unpaired.fastq

# Define variables for second script.

trim1=$(ls *trim1*)
trim2=$(ls *trim2*)

# Second script for filtering PCR duplicates (FilterPCRdupl).

echo ""
echo "======Starting FilterPCRdupl for $pref..."
echo ""

perl $HOME/anaconda3/envs/assembly/bin/filterPCRdupl.pl -fastq1=$trim1 -fastq2=$trim2
    -prefix=$pref -cmp=50

echo ""
echo "======FilterPCRdupl finished for $pref!"
echo ""

# Trimmed reads are zipped and moved to the Trim_reads folder. Useless files are removed.

rm *hist
gzip -r *trim*
mv *trim* ../Output/Trim_reads

# Third script for assembling the reads (SPAdes). Run it and remove the useless files.

echo ""
echo "======Starting SPAdes assembly of $pref..."
echo ""

python $HOME/anaconda3/envs/assembly/bin/spades.py -t 12 --careful --only-assembler -1
    *uniq1.fastq -2 *uniq2.fastq -o ./$pref

rm *uniq*

echo ""
echo "======SPAdes assembly of $pref is finished!"
echo ""

# Go into the folder created by the script, extract 2 files (scaffolds and contigs),
    remove the folder with all the other files and move the two files in the Output folder.

echo "======Compressing reads $pref and moving files..."
cd $pref
mv contigs.fasta $pref.fasta
mv $pref.fasta ../../Output/Contigs
mv scaffolds.fasta $pref.fasta
mv $pref.fasta ../../Output/Scaffolds && cd ../ && rm -r $pref

# rezip the input files and move them back to the original folder.

gzip -r *.fastq && mv ./* ../
cd ../

done < list

rm list
```

#### Quality Control for Genome Assembly Pipeline

This pipeline was designed to run QUAST and to evaluate the overall quality of the genome assemblies (Fig. S1 and Fig. S2). When the values for GC% or total number of contigs is higher than the mean plus twice the standard deviation for the dataset, the assembly is moved to a different folder for further inspection and its name is appended to a list of comparatively low-quality files. During these analyses it was noticed that bovine-specific lineage CC61 tends to have a significantly higher number of contigs than other lineages, when sequencing with short-read technologies and assembling *de novo* (Fig. S2). This CC is known for its high number of mobile genetic elements (MGE) [1,2], which contain repetitive sequences that contribute to genome fragmentation.

---

```
#!/bin/sh

# Run the script from the directory containing the assembled genomes.
# Create directories where to store the output files.

mkdir ./quast_output
mkdir ./quast_output/lowQS

# Run QUAST

python $HOME/anaconda3/envs/assembly/bin/quast -o ./quast_output *.fasta

# Take QUAST output transposed_report.tsv (tab separated with columns = parameters, rows =
    genomes). Remove the first row (header).

sed '1d' ./quast_output/transposed_report.tsv > ./transposed_report_no_header.tsv

# Define variables and calculate the mean values for GC% and number of contigs.

tot_genomes=$(wc -l transposed_report_no_header.tsv | awk '{print $1}')
sum_contigs=$(awk '{s+=$14}END{print s}' transposed_report_no_header.tsv)
sum_GC=$(awk '{s+=$18}END{print s}' transposed_report_no_header.tsv)

mean_contigs=$(echo "scale=2; $sum_contigs / $tot_genomes" | bc)
mean_GC=$(echo "scale=2; $sum_GC / $tot_genomes" | bc)

# Define variables and calculate standard deviation (SD) and 2*SD (SD2).

SD_contigs=$(awk '{sum+=$14; sumsq+=$14*$14}END{print sqrt(sumsq/NR - (sum/NR)**2)}'
    transposed_report_no_header.tsv)
SD_GC=$(awk '{sum+=$18; sumsq+=$18*$18}END{print sqrt(sumsq/NR - (sum/NR)**2)}'
    transposed_report_no_header.tsv)

SD2_contigs=$(echo "$SD_contigs * 2" | bc)
SD2_GC=$(echo "$SD_GC * 2" | bc )

# Define variables and calculate the mean+2SD for contigs and GC.

sum_SD2_contigs=$(echo "$mean_contigs + $SD2_contigs" | bc)
sum_SD2_GC=$(echo "$mean_GC + $SD2_GC" | bc)

rounded_sum_SD2_contigs=$(printf "%.0f\n" "$sum_SD2_contigs")
rounded_sum_SD2_GC=$(printf "%.2f\n" "$sum_SD2_GC")

# If the total number of contigs or the GC% is > than 2SD, print the name of the sequences
    on a list.

awk '{if($14>'$rounded_sum_SD2_contigs' || $18>'$rounded_sum_SD2_GC')print$1".fasta"}'
    transposed_report_no_header.tsv > list_lowQS.txt

# Remove unnecessary files
```

```
rm transposed_report_no_header.tsv
sed 's/_/#/2' list_lowQS.txt >list_names.txt

# Move low-quality score sequences from the list to another folder for further inspection.

for file in $(<list_names.txt);
do
    mv "$file" ./quast_output/lowQS;
done
```

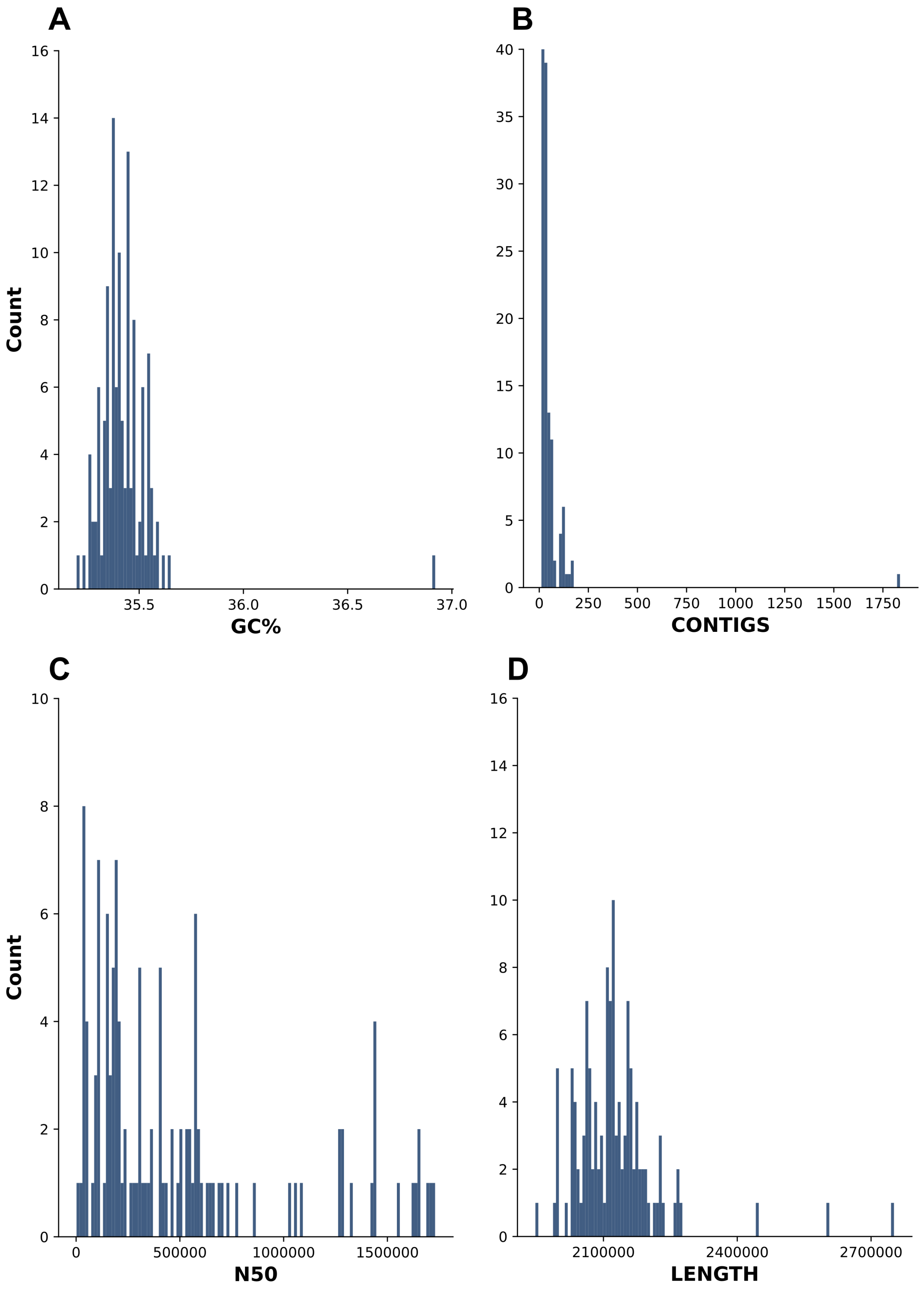


**Figure S1.** Distribution, for the 122 genomes initially included in this study, of GC content (A), total number of contigs (B), N50 (C) and total genome length (D). Two genomes were excluded from the analyses: genome MRI Z2-151 and MRI Z2-309. The latter had a higher GC content compared to the rest of the dataset (A): 36.92%, with a dataset mean value of 35.42% $\pm$0.32 2SD, and was identified as *Enterococcus thailandicus* based on KmerFinder [3]. The former had a higher number of contigs (n = 1,837) (B) and total genome length (2,751,323 bp) (D) compared to the rest of the dataset, which probably indicates contamination.

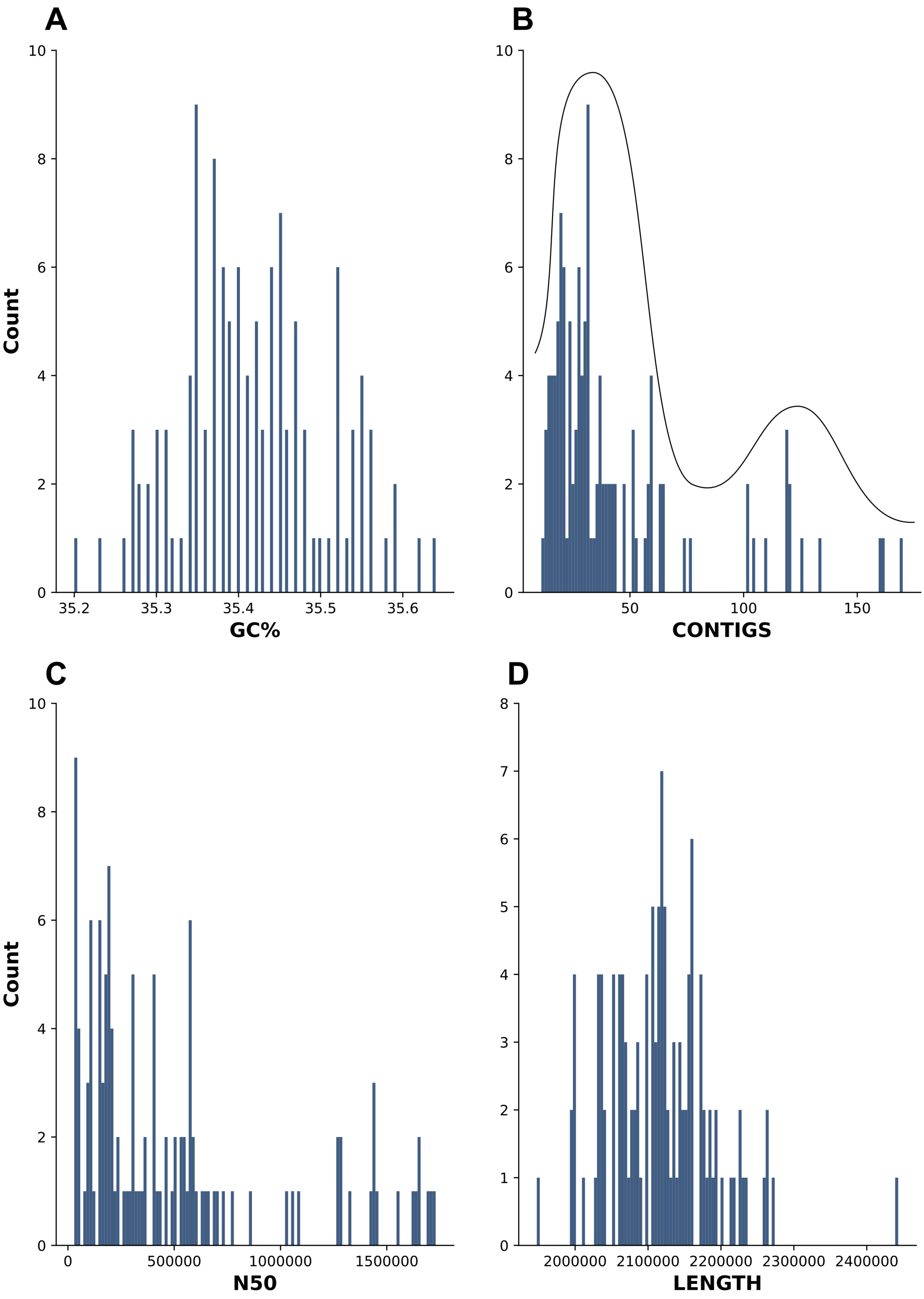


**Figure S2.** Distribution, for 120 high quality group B *Streptococcus* genomes included in the analyses, of GC content (A), total number of contigs (B), N50 (C) and total genome length (D). Panel B shows a bi-modal distribution. This is probably caused by differences in the number of mobile genetic elements, in particular insertion sequences (IS), which incorporate repetitive motifs that impede assembly of short read sequences. Most isolates contributing to the right hand mode belong to bovine-adapted clonal complex 61 (n = 11).

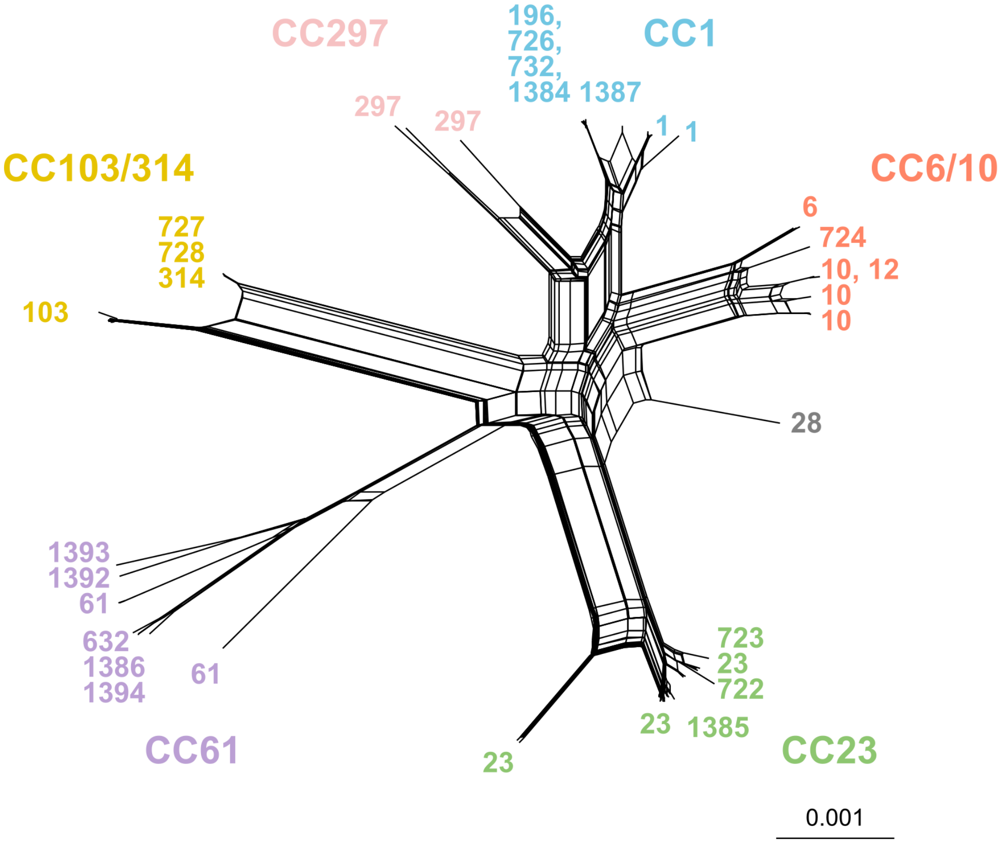


**Figure S3.** Network phylogeny of 120 group B *Streptococcus* (GBS) isolates from bovine milk, collected from 1953 to 1978 (44 historical isolates) and 1997 to 2012 (76 contemporary isolates) shows presence of 6 clonal complexes (CC). Putative recombination is shown as estimated by SplitsTree [4]. Numbers on the leaves represent sequence types (ST).

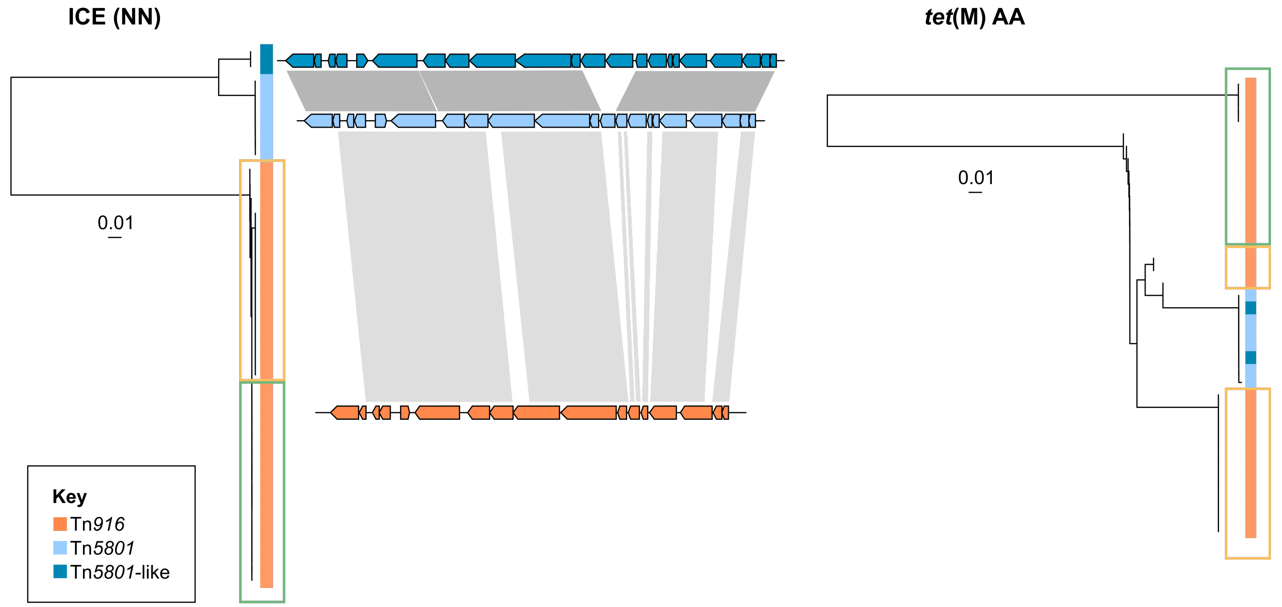


**Figure S4.** Comparative phylogenies of nucleotide sequences (NN) of integrative conjugative elements (ICE) (left) and of the *tet*(M) amino acid sequences (AA, right) encoded by those ICE. Among the genomes of 120 group B *Streptococcus* (GBS) isolates from bovine milk, 37 whole ICE were identified, with gene compositions as visualised with Easyfig v2.2.2 [5]. For sequences derived from Tn*916*, coloured boxes (yellow, green) indicate the distribution of *tet*(M) sequences relative to their ICE sequences. The *tet*(M) gene tree shows clustering of sequences contained by Tn*5801* (n = 6) or Tn*5801*-like (n = 2) ICE within those contained by Tn*916* (n = 29). Tn*916* and Tn*5801* sequences all met the thresholds set for query coverage (QC>80%) and identity (I> 95%), whereas Tn*5801*-like sequences showed high ID% to Tn*5801*, but only partial QC (ID = 99.99%, QC = 93% to the reference). This was due to the substitution of one hypothetical gene with a different hypothetical gene and one IS*256* family transposon in the Tn*5801*-like element, which is observable in the Easyfig comparison.

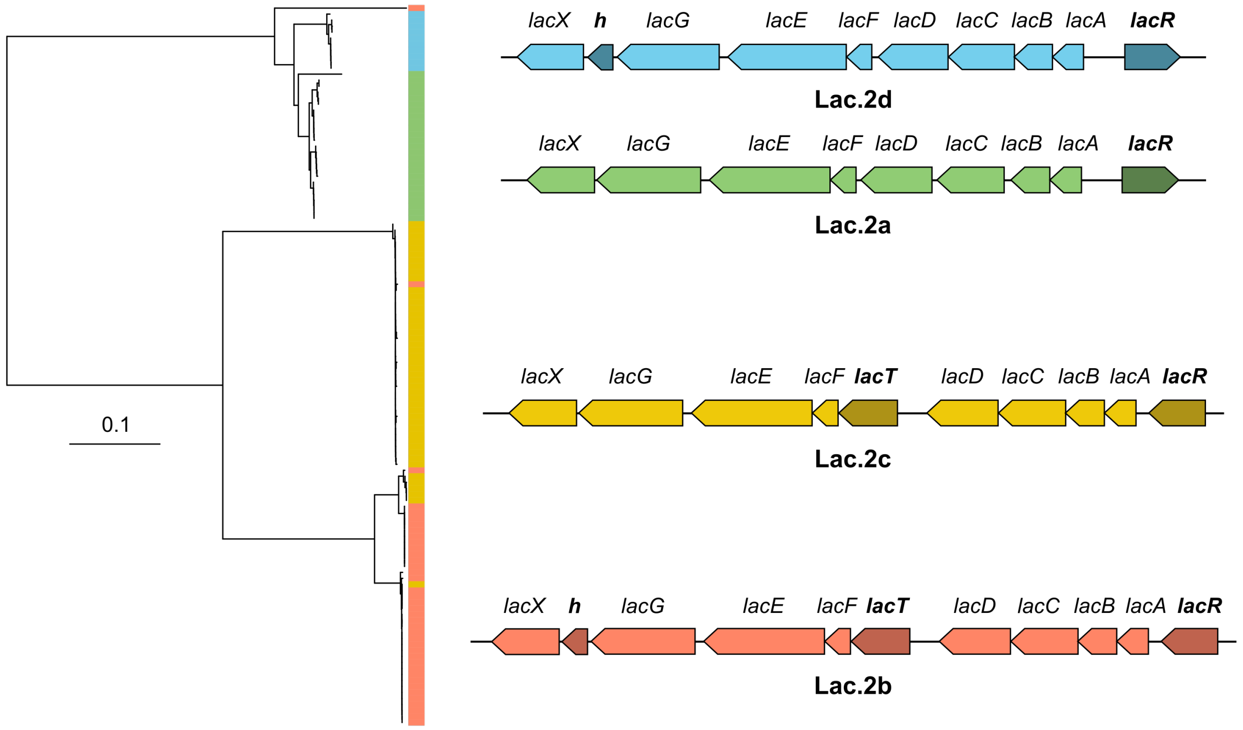


**Figure S5.** Lactose operon Lac.2 was detected in 118 of 120 bovine group B *Streptococcus* (GBS) isolates and could be classified into four variants based on orientation of *lacR* and presence of *lacT* and hypothetical gene (*h*). Gene organisation of Lac.2a, Lac.2b, Lac.2c [6] and Lac.2d (this study) is not consistently concordant with the phylogeny of the operon, indicating gene losses or acquisitions and rearrangements in phylogenetically distant variants, exemplifying the plasticity of the accessory genome of GBS. Phylogeny is based on Lac.2 nucleotide sequences extracted with ARIBA [7] and manually curated. Alignment was obtained with MAFFT [8] and a neighbor-joining tree was constructed with the Jukes-Cantor model in Geneious [9]. Coloured bar next to phylogeny indicates the lac operon variant as shown on the right.
